# Bursts of genomic instability potentiate phenotypic and genomic diversification in *Saccharomyces cerevisiae*

**DOI:** 10.1101/2022.03.24.485697

**Authors:** Lydia R. Heasley, Juan Lucas Argueso

## Abstract

How microbial cells leverage their phenotypic potential to survive in a changing environment is a complex biological problem, with important implications for pathogenesis and species evolution. Stochastic phenotype switching, a particularly fascinating adaptive approach observed in numerous species across the tree of life, introduces phenotypic diversity into a population through mechanisms which have remained difficult to define. Here we describe our investigations into the mechanistic basis of colony morphology phenotype switching which occurs in populations of a pathogenic isolate of *Saccharomyces cerevisiae*, YJM311. We observed that clonal populations of YJM311 cells produce variant colonies that display altered morphologies and, using whole genome sequence analysis, discovered that these variant clones harbored an exceptional collection of karyotypes newly altered by *de novo* structural genomic variations (SVs). Overall, our analyses indicate that copy number alterations, more often than changes in allelic identity, provide the causative basis of this phenotypic variation. Individual variants carried between 1 and 16 *de novo* copy number variations, most of which were whole chromosomal aneuploidies. Notably, the inherent stability of the diploid YJM311 genome is comparable to that of domesticated laboratory strains, indicating that the collections of SVs harbored by variant clones did not arise by a chronic chromosomal instability (CIN) mechanism. Rather, variant clones acquired these complex karyotypic configurations simultaneously, during stochastic and transient episodes of punctuated systemic genomic instability (PSGI). Surprisingly, we found that the majority of these highly altered variant karyotypes were propagated with perfect fidelity in long-term passaging experiments, demonstrating that high aneuploidy burdens can often be conducive with prolonged genomic instability. Together, our results demonstrate that colony morphology switching in YJM311 is driven by a stochastic process in which genome stability and plasticity are integrally coupled to phenotypic heterogeneity. Consequently, this system simultaneously introduces both phenotypic and genomic variation into a population of cells, which can, in turn perpetuate population diversity for many generations thereafter.

## Introduction

The ability to adapt to fluctuating environmental conditions is critical to the survival of organisms. Many adaptation strategies rely on the regulated activities of environmental response pathways, which enable a cell to sense an environmental change, and induce a programmed response such as to alter its phenotype accordingly^1^. Alternatively, cells may stochastically shift between phenotypic states, even in the absence of environmental stimuli. This process, called phenotype switching, generates phenotypic heterogeneity within a clonal population, and can produce subpopulations of cells better equipped to withstand an acute fluctuation in environmental conditions^2,3^. For this reason, stochastic phenotype switching has been proposed to function as a bet hedging strategy, as switching has been found to increase the overall adaptive potential of a population^3-6^.

Stochastic variations in microbial colony morphology (CM) are iconic examples of phenotype switches. Among the best described are the switches displayed by the opportunistic pathogen *Candida albicans*^7,8^. *C. albicans* cells normally form smooth colonies, yet can stochastically switch to form a repertoire of highly differentiated colony structures^8^. Such alterations in colony structure are thought to support population survival in fluctuating environments by enabling cell specialization, resource sharing, and cooperative growth^9,10^. Similar morphological switches have also been observed in the pathogen *Cryptococcus neoformans* as well as in pathogenic isolates of the budding yeast *Saccharomyces cerevisiae*^11,12^. In the context of pathogenesis, the morphological attributes of variant colony architectures are likely to be clinically relevant, as they promote invasive growth, substrate adherence, and biofilm formation^13^. Indeed, switching to a complex morphology has been shown to promote virulence and drug resistance in *C. albicans*^8,14^. This association with virulence, combined with the numerous examples of CM switches among pathogenic microbes^7,15^, underscores the importance of understanding how these switches operate, and how they contribute to the adaptive potential of microbial populations.

CM switching has been described for numerous species, yet the biological mechanisms which underlie these switches remain poorly understood^4^. Since genomic mutations usually impart permanent and heritable genotypic changes to a cell, they have been generally dismissed as putative mechanisms of switching. However, some mutations, such as whole chromosome copy number alterations (CCNAs)(*e.g*., aneuploidies) represent a potentially reversible class of structural genomic variation (SV), as cells can toggle between euploid and aneuploid states simply through stochastic errors in mitotic chromosome segregation^5,16-18^. Indeed, although often portrayed as an exclusively detrimental genomic alteration^19-21^, aneuploidization is also recognized as an adaptive strategy used by cells to survive fluctuating environmental conditions^18,22,23^. Aneuploidies can confer short- and long-term advantages to a cell by acutely and reversibly impacting gene expression and by enabling the subsequent acquisition of permanent adaptive mutations^17,24,25^.

In fact, aneuploidy-driven phenotypic switching has recently been described in the literature, specifically in studies reporting the colony morphology switching behaviors displayed by a laboratory-derived haploid strain of *S. cerevisiae* known as F45^5,26^. In those studies, the authors observed that F45 produced colonies displaying variant morphologies and demonstrated that *de novo* CCNAs were the mechanistic basis of this phenotypic heterogeneity. These studies provided early support for the notion that genotypic mechanisms, specifically aneuploidies, can underlie phenotype switching behaviors. They also raised important and unanswered questions about how the structural stability of the genome can impact the phenotypic plasticity and evolution of microbial communities over time. Here, we investigated the stochastic colony morphology phenotype switching behavior displayed by a naturally derived pathogenic isolate of *S. cerevisiae* YJM311. In the course of defining the complete structural genomic architecture of YJM311^27^, we observed that cells derived from clonal populations of this strain formed variant colony morphologies even when grown in identical conditions. Our subsequent characterization of the genomic architectures of these variant colonies revealed that each had acquired *de novo* structural genomic variations (SVs), and most harbored karyotypes extensively and uniquely altered by *de novo* CCNAs. We used this collection of variant clones to explore the temporal and phenotypic attributes of stochastic aneuploidization in the context of a naturally derived strain, as well as to directly assess the long-term karyotypic stability displayed by these phenotypic variants. Collectively, our results demonstrate that stochastic aneuploidization is a potent driver of genomic diversification within a population and that the long-term stability of the karyotypic alterations derived from these events further perpetuates both phenotypic and genotypic evolution.

## Results

### Phenotypic variants spontaneously arise in clonal populations of the natural isolate YJM311

YJM311 is a diploid clinical isolate of *S. cerevisiae* originally characterized by McCusker and colleagues^28^. While working with this strain for a separate study^27^, we observed that while most cells formed smooth colonies on glucose-rich media (2.0% glucose; YP^2.0%^), ∼1 in 1000 colonies developed a variant complex morphology (average frequency of appearance, 1.29×10^−3^). This variant morphology was evident early in colony formation and became progressively more distinguishable as colonies expanded (Fig. 1A). Like other clinical isolates, YJM311 is known to express several pathogenically relevant phenotypes including the ability to form biofilm-like colonies in response to environmental stimuli, such as changes in the availability of carbon^10,28-30^. Wild type (WT) YJM311 cells grown on YP^2.0%^ normally form smooth colonies (Fig. 1A, top; Fig. 1B, top), but when grown in glucose-poor conditions (0.5% glucose, YP^0.5%^), cells activate a developmental program called the colony morphology response^30^, and form colonies with a complex architecture (Fig. 1B, bottom).

**Figure 1.**
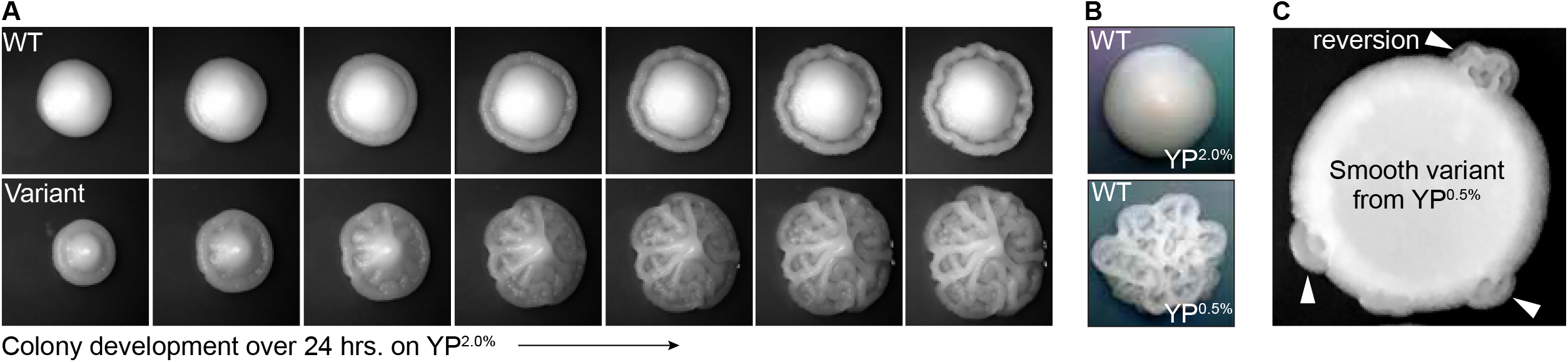
Phenotypic variants spontaneously arise in populations of YJM311. **A**. Representative time-course images of 2-day old WT and complex variant colonies developing over 24 hours on YP^2.0%^. **B**. Representative images of WT YJM311 colony morphologies on YP^2.0%^ (top) YP^0.5%^ (bottom). **C**. An image depicting a cell spot of a smooth variant (6^*sv*^) isolated from YP^0.5%^; the smooth variant phenotype persists throughout the spot, but regions denoted by white arrowheads have reverted to the WT morphology (*i.e*., complex).

The variant complex colonies that appeared on YP^2.0%^ plates were remarkably similar in morphology to YJM311 colonies grown in glucose-poor conditions, suggesting that these cells may have inappropriately activated the colony morphology response, even in the absence of any environmental stimulus. We investigated whether morphological variants also arose among populations of cells plated to YP^0.5%^ media, a condition in which cells normally form complex colonies. Indeed, when we screened colonies grown on YP^0.5%^ plates, we identified smooth morphological variants, also at a frequency of ∼1 in 1000 colonies (average variant frequency, 1.05×10^−3^). We verified that both smooth and complex variants did not form due to local changes in glucose concentration by resuspending these variant colonies in water and spotting them onto new YP^2.0%^ or YP^0.5%^ plates. In all cases, the spotted populations retained the variant phenotype, demonstrating that the variant state was inherent to each clone, and not to the environmental conditions in which they had developed. However, after extended growth, some spots developed sectors which had reverted to the wild-type (WT) phenotype (Fig. 1C, arrowheads), indicating that this phenotypic variation was, in some cases, a reversible phenomenon.

### Phenotypic variants harbor elaborate collections of de novo structural variations

Tan and colleagues previously found that a similar phenomenon of colony morphology variation exhibited by the laboratory haploid strain of *S. cerevisiae* F45 was driven by new mutations, specifically the acquisition of *de novo* chromosomal copy number alterations (CCNAs)^5^. Because the colony morphology phenotypes displayed by YJM311 variant clones appeared to be heritable over many generations (Fig. 1C), we considered the possibility that a genetic mechanism might also underlie the phenotypic heterogeneity displayed by this strain and investigated this hypothesis by performing comprehensive whole genome sequencing (WGS) analysis on collections of clones which displayed either the WT morphology (20 clones), or complex (22 clones isolated from YP^2.0%^) and smooth (26 clones isolated from YP^0.5%^) variant morphologies, respectively.

We expected that all WT clones would harbor the parental YJM311 karyotype, yet, our WGS analysis demonstrated that 20.0% (4/20) were monosomic for chromosome 1 (Chr1). Our recent genomic characterization of this strain revealed that one homolog of Chr1 (Chr1*b*) is a ring chromosome^27^. Analysis of WGS data generated from the parental YJM311 stock established that it is was a mosaic population consisting of euploid cells and cells lacking Chr1*b* (Table S1). Indeed, Chr1*b* was the absent homolog in all four monosomic WT clones, suggesting that these clones were derived from monosomic cells pre-existing in the stock population. Among the collection of complex and smooth variant clones, 9.5% (2/21) and 15.3% (4/26) lacked Chr1*b*, respectively (Table S2). Because cells lacking Chr1*b* can form colonies displaying the WT morphology on both YP^2.0%^ and YP^0.5%^ plates, monosomy of Chr1 is unlikely to be linked to the phenotypic variation we were investigating. For this reason, we excluded aneuploidy of Chr1*b* from our subsequent genomic analyses of WT and variant clones.

Besides aneuploidy of Chr1*b*, we detected no other karyotypic alterations in the genomes of the set of WT clones. In striking contrast, however, we detected other *de novo* structural variations (SVs) in the genomes of every single variant clone (Fig. 2A, D, E). Intriguingly, the majority of these new SVs were CCNAs (89%), and most often constituted chromosomal gains (Fig. 2B), a pattern consistent with that spectra reported by Tan *et al*. for morphological variants derived from the strain F45^5^. Several variant clones harbored non-reciprocal translocations which resulted in segmental copy number variations of specific chromosomal regions (4/47 variant clones) (Fig. 2B, Seg. CNVs; Fig. S1; Table S2). In all cases, these chromosomal rearrangements had boundaries at Ty elements, suggesting that they likely arose through non-allelic homologous recombination between dispersed repeats (Fig. S1, Table S2). We also detected several copy-neutral tracts of loss of heterozygosity (CN-LOH) in variant genomes that had also been altered by CCNAs (Fig. 2B, CN-LOH; Table S2). Collectively, these results suggested that *de novo* copy number variations, especially in the form of CCNAs, were likely the cause underlying the colony morphology variation displayed by YJM311.

**Figure 2.**
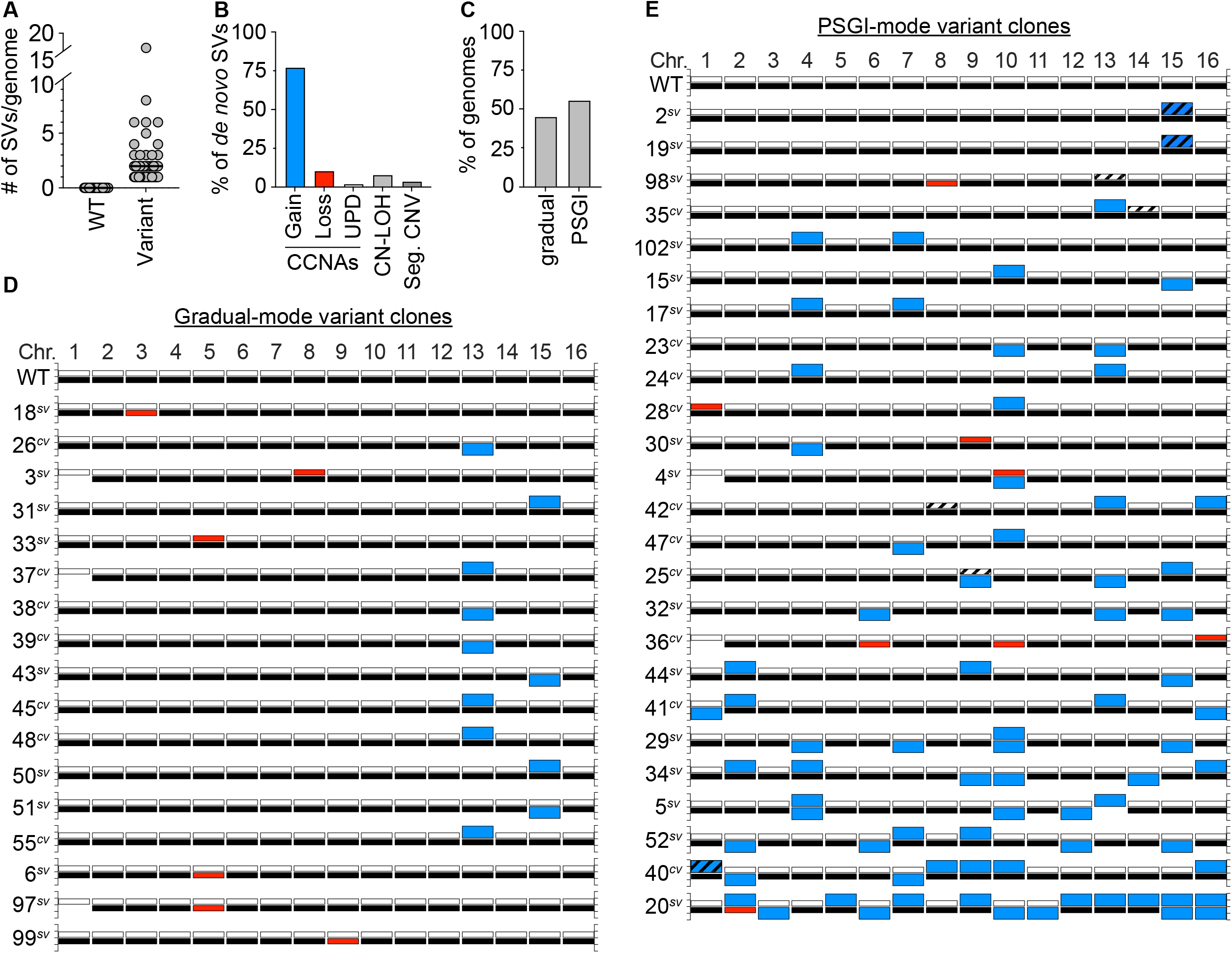
Variant clones harbor numerous *de novo* SVs. **A**. The number of SVs present in WT and variant genomes. Each grey circle denotes an individual clone. Black bars denote the median number of SVs for each set. **B**. The percentage of SVs that constitute whole chromosomal gain, loss, uniparental disomy (UPD), copy-neutral loss-of-heterozygosity (CN-LOH), or segmental copy number variations (Seg. CNV). **C**. The percentage of variant genomes displaying either a gradual or PSGI mode of mutagenesis. **D-E**. Karyotypes of the YJM311 parent and all variants arising by gradual and PSGI modes of aneuploidization. For each chromosome, white bars represent homolog ‘*a*’ and black bars represent homolog ‘*b*’. Blue bars denote chromosomal gains, red bars denote chromosomal losses. Striped bars denote tracts of CN-LOH. Loss of Chr1*b* has been excluded from analysis in A, B, and C.

Overall, the degree to which these variant genomes had been impacted by SVs fell along a spectrum, with individual clones harboring between 1 and 16 SVs per genome (median, 2 SVs/ genome) (Fig. 2A, D, E; Table S2). The most extremely altered variant clone in our collection of variants was the smooth variant 20 ^*sv*^, which had acquired the following CCNAs: uniparental disomy (UPD) of Chr2 (a type of CCNA that results in copy-neutral whole chromosome LOH through concurrent loss of one homolog and gain of the other^16,31^), trisomy of Chr3, Chr5, Chr6, Chr7, Chr9, Chr10, Chr11, Chr12, Chr13 and Chr14, and tetrasomy of Chr15 and Chr16 (Fig. 2E, Table S2). All the SVs harbored by 20^*sv*^, as well as those harbored by all other clones, were detected at read-depth coverages consistent with discrete integer copy number levels, suggesting that they were acquired simultaneously, and did not arise continuously during colony expansion (data not shown). Given that all variant clones were isolated from populations that had been limited to only ∼30 generations of growth, this spectrum of karyotypic alterations prompted us to further characterize the mutational patterns underlying the apparent genomic evolution which had occurred in these clones. Recent work from our group and others has demonstrated that stochastic acquisition of SVs occurs primarily through two distinct modes: 1) the well-established neo-Darwinian pattern of gradual mutation accumulation, during which single SVs are acquired independently over time, and 2) a burst-like pattern characterized by transient episodes of punctuated systemic genomic instability (PSGI), during which multiple SVs are acquired simultaneously^32-36^. Our WGS analysis indicated that 44.6% of clones had acquired just a single *de novo* SV and best fit a gradual mode of mutagenesis (Fig. 2C). However, the majority of variant clones (55.4%) had acquired two or more *de novo* SVs in the same time frame, and better fit a PSGI mode of mutation accumulation (Fig. 2C)^32^.

Most variants harbored unique karyotypic configurations (Fig. 2D, E; Table S2). However, among the variant clones which displayed a gradual pattern of aneuploidization, we identified an enrichment for variants which had become trisomic for either Chr13 or Chr15 (Fig. 2D; Fig. S2A). The recovery of such clones indicated that while not necessary, trisomy of Chr13 or Chr15 is sufficient to induce detectable variations in colony morphology in YJM311. Allelic differences between genomic loci present on Chr13 and Chr15 are known to modulate colony morphology phenotypes and expression of the colony morphology response in YJM311^30^, yet, we observed no bias for the gain of specific homologs of Chr13 or Chr15 in our set of variant clones. Moreover, variant clones harboring extra copies of the different homologs of either Chr13 or Chr15 only modestly differed from one another in colony morphologies, respectively (Fig. S2B, C). Taken together, these results support our conclusion that *de novo* SVs resulting in copy number variations, not copy-neutral SVs resulting only in changes in allelic identity, underlie the stochastic appearance of morphological variants in populations of YJM311.

### YJM311 cells do not display elevated levels of chromosomal instability

The patterns of SVs harbored by most variant clones closely matched those defined in previous studies of PSGI^32,35^. However, because aneuploid variants arose relatively frequently in populations of YJM311 cells, we explored the possibility that YJM311 might inherently display the mutator phenotype known as chromosomal instability (CIN), which is characterized by increased rates of aneuploidization and chromosomal rearrangements^37-39^. To determine if YJM311 cells acquired CCNAs at elevated frequencies, we used a counter-selectable chromosome loss assay and fluctuation analysis to calculate the rate at which cells lost Chr5*b*^27,32,40^. Using a strategy similar to that described in our previous study dissecting the mutational spectra of PSGI aneuploidization events^32^, we deleted the counter-selectable marker *CAN1* from its endogenous locus on the left arm of Chr5*a* and introduced a second copy of *CAN1* onto the right arm of Chr5*b* (Fig. S3A)^32^. Due to the presence of the two copies of the *CAN1* gene on Chr5*b*, this diploid parental strain was sensitive to the toxic arginine analog canavanine. However, cells that lost Chr5*b* acquired resistance to the drug, and could form colonies on media containing canavanine^40^. The average rate at which YJM311 cells lost Chr5*b* was 1.04×10^−5^/cell/generation (Table S3), a value consistent with our published rates of chromosome loss in several well-defined laboratory *S. cerevisiae* strains^32,41-43^. For the purposes of comparison, we also performed this same assay using a well-defined and karyotypically stable diploid laboratory hybrid strain (JAY3106), which had been constructed in a manner identical to the strategy used for YJM311. This strain lost the *CAN1*-marked homolog of Chr5 at a rate nearly identical to that of YJM311 (1.03×10^−5^/cell/generation)(Fig. S3; Table S3). Because YJM311 did not show elevated rates of chromosome loss relative to these other strains, we conclude that a chronic CIN phenotype is unlikely to drive the pervasive aneuploidy we observed in the genomes of variant YJM311 colonies. Rather, the frequent appearance of aneuploid variants in populations of YJM311 is more likely to be a combined consequence of the contributions of multiple genomic loci to the expression of the colony morphology response^30^, and the finding that PSGI events underlie the formation of most phenotypic variants. Because PSGI-derived variants harbor multiple CCNAs, each CCNA may differentially impact the expression of one or more response-linked loci.

### The stability of variant karyotypes impacts long-term genomic and phenotypic diversification

Aneuploidy has been shown to impart pleiotropic negative effects to the growth and genomic integrity of yeast cells, presumably due to imbalances in gene expression and dosage^44,45^. Indeed, previous studies have demonstrated that haploid cells forced to carry extra copies of different chromosomes display distinct signatures of genomic instability, including varying severities of CIN^44^. These and other studies have supported a model positing that aneuploidies are sufficient impart a chronic CIN phenotype to cells^38^. Yet, our WGS analysis suggested that the aneuploid karyotypes harbored by YJM311 phenotypic variants were conducive with prolonged genomic stability, as they appeared to be stably propagated throughout the growth of the variant colony. This was surprising, particularly for the variant clones which had acquired the highest burdens of aneuploidies through PSGI-type events (Fig. 2E; *e.g*., 20^*sv*^, 40^*cv*^).

Recent work has demonstrated that some wild isolates of *S. cerevisiae* display enhanced aneuploidy tolerance relative to those derived from a laboratory strain called w303a^44,45^. w303a has been used extensively in studies exploring the physiological consequences of aneuploidization and was recently shown to be particularly sensitive to the gene dosage effects imparted by whole chromosome gains^44,46^. These recent studies linked the aneuploidy sensitivity displayed by w303a to single nucleotide polymorphisms in the gene encoding the translational regulator *SSD1*^46^. Because the aneuploid variant colonies recovered from YJM311 appeared to stably propagate *de novo* CCNAs and other CNVs, we predicted that YJM311 may harbor copies of *SSD1* lacking the sensitizing *SSD1* allele present in w303a. To determine if YJM311 carried the known sensitizing allele of *SSD1*, we interrogated the available phased diploid genome assembly of YJM311 and found that neither of the *SSD1* alleles encode the polymorphisms posited to confer aneuploidy sensitivity to w303a (data not shown)^27^. Thus, the short-term karyotypic stability inferred from our WGS analysis of aneuploid variant clones could reflect general aneuploidy tolerance inherent to YJM311.

We wished to more directly evaluate the long-term stability of the aneuploid karyotypes carried by our collection of aneuploid phenotypic variants, as we were interested in determining whether these karyotypes would impart chronic CIN phenotypes to variant cells and whether overall aneuploidy burden is a predictor of karyotypic and phenotypic stability. To investigate these questions, we performed mutation accumulation (MA) experiments in which we passaged 31 variant clones and 9 WT euploid controls for ∼250 generations through single colony bottlenecks on solid media (Fig. 3A). We performed this experiment for variant clones harboring karyotypic configurations that had resulted from both gradual (8 clones) and PSGI modes (23 clones) of aneuploidization. At every passage, we preserved a stock of each single-colony bottleneck, and after completion of the experiment, we analyzed the genomes of the terminal MA lines using WGS analysis and compared them to the genomes of the parental variant clones.

**Figure 3.**
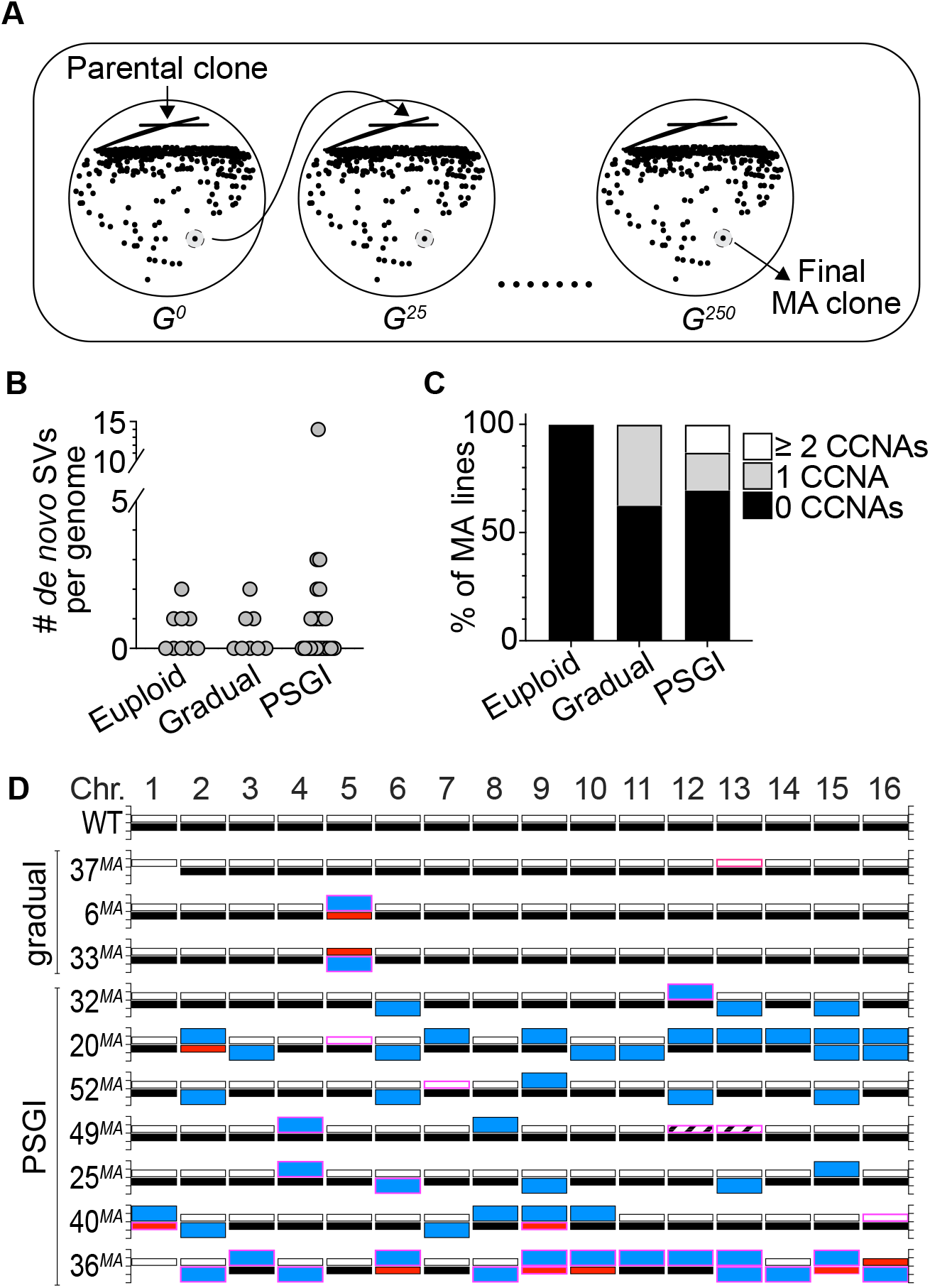
Phenotypic variants display long-term genomic stability. **A**. A cartoon schematic outlining the mutation accumulation study performed on euploid and variant clones. **B**. The number of *de novo* SVs harbored by the delineated subsets of terminal MA lines. **C**. The percentage of MA lines harboring the denoted numbers of *de novo* CCNAs. For each subset, median number of *de novo* SVs equals zero. **D**. Karyotypes of the MA lines derived from gradual and PSGI-type variant clones that acquired *de novo* CCNAs over the course of the experiment. Bars outlined in pink denote *de novo* CCNAs that arose during the MA study. Bar colors same as in Fig. 2.

To establish the basal level of *de novo* genomic change that occurred over the course of 250 generations of growth, we first characterized the genomes of the euploid control MA lines and found that four had acquired *de novo* tracts of CN-LOH (Fig. 3B, Table S4). This was expected, as normal mitotic recombination processes active during vegetative propagation can result in CN-LOH^47^. The pooled rate at which CN-LOH occurred in the genomes of the euploid MA lines was similar to rates reported from similar MA studies (2.2×10^−3^/division)^47,48^. This, together with our finding that the euploid MA lines harbored no other *de novo* genomic alterations after 250 generations (Fig. 3C), lends additional support to our above conclusion that the YJM311 genome is effectively stable (Fig. S3).

Next, we interrogated the genomes of the MA lines derived from the gradual-mode variant clones and found that most karyotypes remained unchanged (Table S4). Only three had acquired *de novo* SVs over the course of the MA experiment and in each case, these three clones (37^*MA*^, 6^*MA*^, and 33^*MA*^) had acquired a *de novo* CCNA (Fig. 3B, 3C, 3D; Table S4). Intriguingly, all three *de novo* CCNAs returned the variant genomes to a state of euploidy, respectively (Fig. 3D). The MA line derived from 37^*cv*^ (37^MA^), which was trisomic for Chr13 at the beginning of the experiment, had lost one copy of Chr13 and was again disomic for all chromosomes (Fig. 3D; Table S4). Likewise, the variant clones 6^*sv*^ and 33^*sv*^ had both been monosomic for Chr5 at the beginning of the experiment, and the MA lines derived from each had acquired a second copy of the remaining homolog of Chr5 and were again disomic (Fig. 3D; Table S4). We detected only one other SV among the MA lines in this subset of gradual-mode variant clones, a CN-LOH event that had occurred between the three copies of Chr13 in 37^*cv*^. Collectively, the median number of *de novo* CCNAs harbored by the MA lines derived from the gradual-mode variants was zero, and the pooled rate at which these clones acquired *de novo* CCNAs was 1.5×10^−3^/cell/div.

We then analyzed the genomes of the MA lines derived from the PSGI-mode variant clones. Because they harbored extensive aneuploidy burdens, we expected that the genomes of these MA clones would show evidence of CIN and continued karyotype evolution. Strikingly, however, our analysis revealed that the majority of PSGI-class variant karyotypes had been stably propagated throughout the duration of the experiment (16/23)(Fig. 3B, 3C; Table S4). Twelve MA lines harbored karyotypes identical to their respective ancestors, and 4 MA lines differed from their ancestral parents only by *de novo* tracts of CN-LOH (Table S4). Only seven variant MA lines had acquired *de novo* CCNAs (Fig. 3C, 3D). Like the gradual-class of MA lines, the median number of *de novo* CCNAs harbored by the combined subset of PSGI-class MA lines was also zero, and the pooled rate at which PSGI-mode MA lines acquired *de novo* CCNAs was only 2.7-fold higher than that of the gradual-class variant clones (4.0×10^−3^/cell/div), demonstrating that numerical aneuploidy burdens are not simple predictors of long-term genomic stability.

Importantly, however, this pooled rate did not accurately represent the collective stability of the PSGI clones, as a single MA line derived from the variant 36^cv^ had acquired 60.9% (14/23) of the total *de novo* CCNAs detected across the entire set of PSGI-mode MA lines. Our earlier genomic analysis of the original complex variant 36^cv^ revealed that it had experienced a PSGI aneuploidization event which resulted in the acquisition of three CCNAs: monosomies of Chr6, Chr10, and Chr16 (Fig. 2E, Table S3). The terminal MA-derived clone, 36^MA^, harbored trisomies of seven chromosomes (Chr2, Chr3, Chr4, Chr8, Chr11, Chr12, and Chr14), tetrasomy of Chr13, and UPD of Chr6, Chr9, Chr10, Chr15, and Chr16. In total, only the homologs of Chr1, Chr5, and Chr7 remained unaffected by aneuploidization. The extensive karyotypic evolution evident in the 36^*MA*^ genome suggested that the original PSGI aneuploidization event that had given rise to the parental variant (36^*cv*^) may have also resulted in the subsequent expression of a CIN phenotype. Indeed, the rate of *de novo* CCNA acquisition calculated from the MA experiment for 36^*cv*^ alone was 34-fold and 37-fold greater than both pooled gradual-mode and PSGI-mode MA lines, respectively (5.6×10^−2^/cell/div). Yet, while the karyotypic evolution of 36^*cv*^ satisfied the well-accepted definition of CIN on the basis of displaying an elevated rate of aneuploidization, we wished to further define the dynamics by which the altered karyotype of 36^*MA*^ had developed over time. If the karyotype harbored by 36^*MA*^ had formed as the result of CIN, then the genomes of the bottleneck clones preserved at each passage in the MA experiment would be expected to reflect chronic instability and the progressive development of the terminal karyotype. Remarkably, however, WGS analysis of the preserved bottleneck clones revealed that 36^*MA*^ had acquired all 14 *de novo* CCNAs within a single passage early in the MA experiment, after which the genome remained stable the next ∼225 generations (Fig. S4). These results demonstrated that the karyotype of 36^*MA*^ had not formed as the result of a CIN phenotype. Rather, 36^*MA*^ appears to have acquired its complex karyotype through a second PSGI event, which was subsequently propagated with perfect fidelity for many generations thereafter (Fig. S4).

Collectively, the results of the MA experiment indicated that most variant karyotypes, even those severely altered by aneuploidization, were conducive with long-term genomic stability. Consequently, the variant morphology phenotypes displayed by these stable clones were also long-lived. However, among the MA lines which had acquired new CCNAs over the course of the MA experiment, we observed that two lines, 6^*MA*^ and 33^*MA*^, had reverted to expression of the WT colony morphology phenotype, and once again formed complex colonies on YP^0.5%^ plates (Fig. 4A). Such an ability to toggle between WT and variant phenotypic states is a defining feature of most switching systems described in the literature, including the previously described aneuploidy-based phenotypic switch exhibited by F45 cells^4,5^. In that study, Tan *et al*. reported that the disomic morphological variants derived from F45 also produced derivatives which had reverted to the WT phenotype, and these reversions were always accompanied by a return to euploidy through a secondary aneuploidization event^5^. Our own results supported this earlier observation, as both 6^*MA*^ and 33^*MA*^ had also returned to euploidy via aneuploidization. The ability of some variant clones to frequently revert to the WT phenotype provoked an interesting premise, namely that the karyotypic stability of a variant could modulate the toggle-like quality of the phenotypic switch. Indeed, cells monosomic for Chr5 readily produced phenotypic revertants, which manifested as the formation of complex sectors in smooth colonies, suggesting that this karyotypic configuration was particularly destabilizing (Fig. 4B, white arrowheads). To determine if the revertant sectors produced by both 6^*sv*^ and 33^*sv*^ had arisen through the same UPD-type mechanism as had occurred in both the 6^*MA*^ and 33^*MA*^ MA lines, we performed molecular karyotype analysis using pulsed field gel electrophoresis (PFGE), a technique that resolves intact yeast chromosomes by size, for the genomes of the parental 6^*sv*^ variant and five revertant sectors which had developed in colonies (Fig. 4C). The two homologs of Chr5 in YJM311 differ from each other by ∼54kb and are particularly well-resolved by PFGE-based molecular karyotype analysis^27^. 6^*sv*^ had lost Chr5*b*, the larger of the two Chr5 homologs^27^ (Fig. 3D; Table S2). Thus, if the formation of revertant sectors was due to acquisition of a second copy of Chr5, then the intensity of the smaller band corresponding to the other homolog of Chr5, Chr5*a*, should increase to a level indicating the presence of two copies. In support of the WGS sequencing analysis of 6^*MA*^ and 33^*MA*^, this PFGE band intensity analysis revealed that all revertant sectors derived from 6^*sv*^ had acquired a second copy of Chr5*a* (Fig. 4D, blue trace). Taken together, these results suggest that in YJM311, the plasticity of a variant phenotype is directly modulated by the stability of the variant genome.

**Figure 4.**
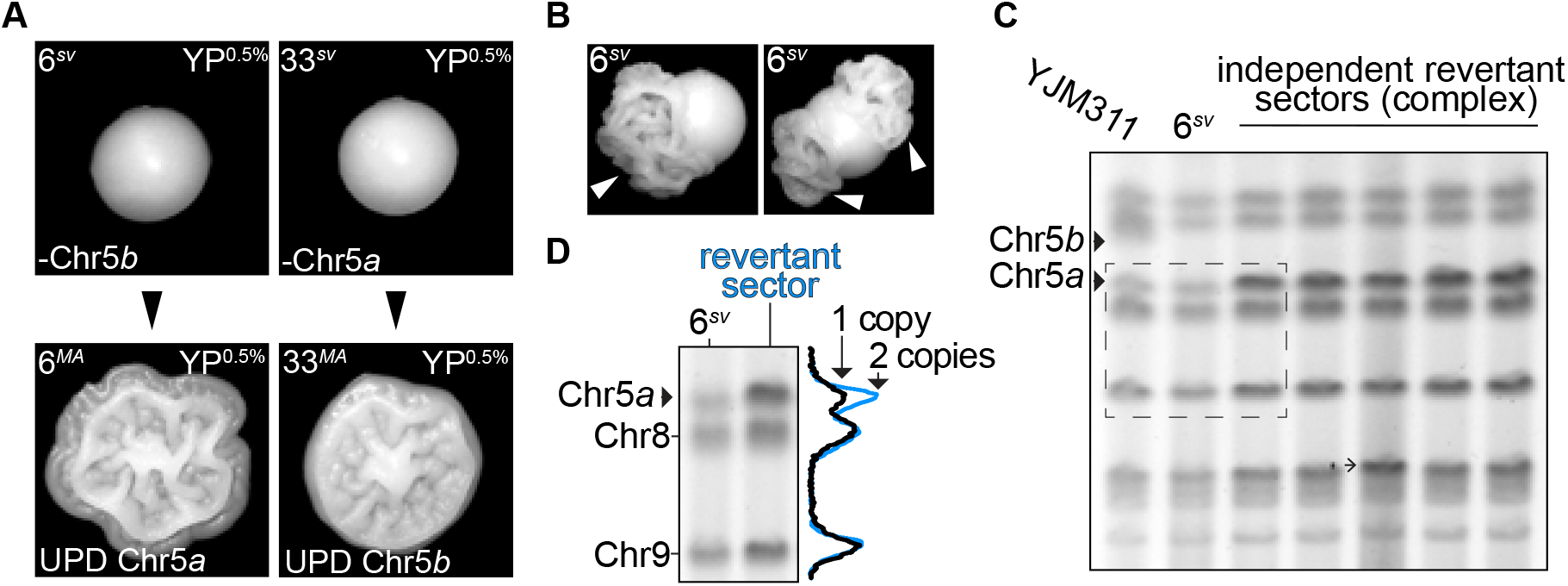
Phenotypic reversions are driven by subsequent aneuploidization events. **A**. Representative images of colony morphologies of variant clones 6^*sv*^ and 33^*sv*^ and their respective MA-derivatives 6^*MA*^ and 33^*MA*^ when grown on YP^0.5%^. **B**. Representative images of sectored colonies derived from 6^*sv*^. **C**. Molecular karyotypes of the parental YJM311 genome, the smooth variant 6^*sv*^, and five revertant sectors recovered from independent colonies. Solid arrowheads denote the bands correlating to the two homologs of Chr5 (Chr5*a*, Chr5*b*). 6^*sv*^ lost Chr5*b*. Barbed arrowheads denote additional SVs that arose concurrently in the genomes of revertant sectors. **D**. Band intensity tracing analysis and representative crop from C. (dashed rectangle) confirming UPD of Chr5*a* in revertant complex sectors. Black trace represents band intensity derived from 6^*sv*^, blue trace represents band intensities derived from complex sectors. The increased amplitude of the blue peak corresponding to Chr5*a* correlates with an increase in copy number from 1 to 2 copies.

## Discussion

Here we have presented data demonstrating that the stochastic colony morphology switching behavior of a pathogenic isolate of *S. cerevisiae* is driven by dynamic structural evolution of the genome. These results directly support and expand upon earlier work by Tan *et al*. describing an aneuploidy-driven colony morphology switch displayed by a haploid strain of yeast^5^, and further substantiate the model that aneuploidy can underlie rapid and reversible phenotype switching events that impact physiological characteristics relevant to the environmental context from which this isolate was recovered (*i.e*., invasive growth, biofilm formation). This observation that stochastic structural genome evolution is coupled with phenotypic plasticity in *S. cerevisiae* contributes an important perspective to our understanding of how phenotypic diversification can shape the genomes of organisms and the diversity of populations.

The potential impacts of aneuploidy-driven phenotype switching on population diversity are further heightened when framed within the emergent concept of PSGI^32-35^. Indeed, the growing body of work substantiating the occurrence of these interesting mutagenic events presents significant implications for our current interpretations of numerous evolutionary processes including microbial adaptation^32-35,49-51^. These events can produce a spectrum of remarkably altered karyotypes^32^, such as those harbored by the phenotypic variants characterized in the present study. Here, our results highlight how the systemic nature of PSGI-type aneuploidization can generate genotypically unique cells that display similarly altered phenotypes (*e.g*., the formation of complex colonies on YP^2.0%^). Perhaps the most interesting implication of aneuploidization as it relates to phenotype diversification derives from our finding that the new karyotypic configurations harbored by stochastically arising phenotypic variants are, for the most part, very stable. Collectively, this stochastic system appears to introduce several types of enduring diversity into a population of microbial cells: 1) diversity in colony morphology and the associated environmental response phenotypes, and 2) karyotypic diversity among these phenotypic variants.

Adaptation through aneuploidization may be especially important for the survival of pathogenic microbes, which must endure fluctuating conditions within their host niches. This is supported by recent genomic surveys which have revealed pervasive aneuploidy in the genomes of pathogenic and clinical isolates of *S. cerevisiae, C. albicans*, and *C. neoformans*^23,52-54^. Moreover, aneuploidy has been associated with increased virulence^8,14^, thermotolerance^17^, and drug resistance^22^ in these fungal pathogens. Intriguingly, these pathogenic traits have also been associated with an isolate’s ability to develop different colony morphologies, be it through a regulated program like the colony morphology response, or through stochastic phenotype switching^8,13-15,55^. Although yet untested, it is tempting to speculate that stochastically arising aneuploid phenotypic variants, such as those characterized in this study, play key roles in pathogenic adaptation and persistence in a disease context. Future studies will focus on defining the contributions of aneuploidy-based phenotypic switches to population survival and adaptation, as well as how these switch events perpetuate genomic evolution and diversity over time.

## Experimental Methods

### Strain Construction

All experiments performed in this study were carried out using the clonal strain of the diploid clinical isolate YJM311, called YJM311-3^27^. To construct the YJM311 strain used for the *CAN1* counterselection assays, a single copy of *CAN1* was deleted from Chr5*a* using a PCR product encoding the *HPHMX6* cassette, which confers resistance to the drug hygromycin B^56^. Correct targeting to and deletion of the *CAN1* locus was confirmed by PCR. Next, a PCR product consisting of *CAN1-KANMX* amplified from genomic DNA was integrated on the right arm of Chr5*b* at position 256375-257958 of the reference genome. Correct insertion or deletion of the *CAN1* gene was confirmed by PCR.

### Variant Recovery and Media

To isolate spontaneously arising complex or smooth phenotypic variants, YJM311 cells were streaked on solid YP^2.0%^ (10g/L yeast extract, 20 g/L Peptone, 20 g/L bacteriological agar, and 20g/L glucose (2.0%)) media and incubated at 30°C for 32 hours to allow single colonies to grow. Individual single colonies were each inoculated into 5mL liquid YP^2.0%^ cultures and incubated at 30°C for another 24 hours on a rotating drum. These cultures consisted of ∼10^8^ cells, or approximately 30 generations derived from the original colony-forming cell. Cultures were diluted appropriately and plated such that ∼200 individual cells were plated to either YP^2.0%^ or YP^0.5%^ (10g/L yeast extract, 20 g/L Peptone, 20 g/L bacteriological agar, and 5g/L glucose (0.5%)). Plates were incubated at 30°C for 3 days and then visually screened for colonies that displayed the variant phenotype. Total colonies were counted and the frequencies at which morphological variants appeared were calculated. The average frequency of the appearance of complex variants on YP^2.0%^ was calculated from a total of 74,155 colonies derived from 48 independent cultures. The average frequency of the appearance of smooth variants on YP^0.5%^ was calculated from a total of 23,342 colonies derived from 22 independent cultures. Candidate variant colonies were isolated, resuspended in water, spotted to fresh plates, and grown at 30°C for at least 3 days to confirm the persistence of the phenotypic variation. The entirety of each patch was then resuspended and frozen until DNA was prepared for genomic sequencing.

### Whole genome sequencing analysis

The genomes of 68 wild-type and variant clones were sequenced using Illumina short read whole genome sequencing and analyzed as described previously^32^. Briefly, Genomic DNA from each clone was isolated using the Yeastar Genomic DNA kit from Zymo Research. Pooled, barcoded libraries of individual genomes were generated using a Seqwell plexWell-96 kit. The final barcoded library was sequenced using an Illumina HiSeq sequencer. Illumina reads for each genome were mapped to the yeast reference genome (R64-2-1, yeastgenome.org) using CLC-Genomics software (Qiagen). Resulting read mapping files were then subjected to copy number and heterozygous single nucleotide polymorphism (hetSNP) variant analysis using the Nexus Copy Number software program (Biodiscovery). From this, we identified whole chromosome gains/losses, segmental duplications/deletions, and tracts of loss-of-heterozygosity (LOH). Aneuploidy was defined as the deviation in copy number of each individual homolog away from 1n. Using this definition, UPDs were scored as two CCNAs. SVs identified in Nexus were confirmed manually using CLC Genomics. The complete karyotypic analyses of variant clones are reported in Table S2. Several clones harboring chromosomal translocations (1^*sv*^, 22^*sv*^, 27^*sv*,^ and 46^cv^) were further analyzed using Oxford Nanopore Technologies (ONT) Minion sequencing. High molecular weight DNA was prepared from cells, barcoded using the ONT ligation sequencing and native barcoding kits (SQK-LSK-109, EXP-NBD-104), and sequenced on a Minion flow cell (FLO-MIN106D). Reads were basecalled using ONT Guppy software and analyzed using CLC Genomics Workbench.

To investigate the mosaicism of Chr1 monosomy in our YJM311 stock and WT clones, we measured the log2 ratio of read coverage across a defined region of Chr1 (position 50,000-150,000). For this analysis, a log2 ratio of ∼0 indicates that the chromosomal region is present at 2 copies, and a log2 ratio of -1 indicates that the chromosomal region is present at only one copy. Log2 ratio values that fall in between 0 and -1 indicate that the sample contains a mixture of cells harboring 1 or 2 copies of the defined genomic region. The log2 values, and corresponding copy number values for Chr1 are reported in Table S2. For comparison, we also measured the log2 ratio of read coverage at region on Chr7 (position 200,000-300,000) within these same genomes. Log2 values and corresponding copy number values for this region are also presented in Table S2 and demonstrate that copy number mosaicism in these clones is restricted to Chr1.

### Fluctuation Analysis

Cells were streaked to YP^2.0%^ and grown at 30°C for 48 hours to allow single colonies to form. Individual colonies were resuspended in 200uL 1x TE buffer (Thermofisher), diluted appropriately, and plated onto YP^2.0%^ and canavanine-supplemented plates (20g/L glucose, 5g/L ammonium sulfate, 1.7g/L yeast nitrogen base without amino acids, 1.4g/L arginine dropout mix, 20g/L bacteriological agar, 0.6mg canavanine sulfate). Plates were incubated at 30°C for 72 hours after which colonies were counted. Colony count data were used to calculate rates and 95% confidence intervals using Flucalc, a MSS-MLE (Ma-Sandri-Sarkar Maximum Likelihood Estimator) calculator for Luria-Delbrück fluctuation analysis (flucalc.ase.tufts.edu)^57^. Calculated rates and confidence intervals are presented in Table S3.

### Mutation accumulation assays

Cells from frozen stocks of the variants and euploid controls denoted in Table S4 were streaked to either YP^0.5%^ or YP^2.0%^ and grown at 30°C for 48 hours. At 48 hours, a single colony from each original plate was picked with a sterile toothpick. Cells from the colony were first re-steaked to a fresh plate and then also resuspended in a glycerol solution and frozen at -80°C. This passaging and preservation procedure was repeated for each clone for 11 more passages, for a total duration of 24 days, and a total generation time of approximately 250 generations (25 generations per 48 hour growth period). To eliminate any bias for specific phenotypic characteristics, colonies selected for passage were selected before colony morphology phenotypes were readily visible. At the end of the experiment, terminal MA clones were sequenced using Illumina sequencing and analyzed as described above.

### Colony Imaging

Colonies were imaged using a Dino-lite Edge 1.3MP AF4115ZT polarized microscope and the companion DinoScope 2.0 software program. Images were processed using Adobe Photoshop, Premier Pro, and Premier Rush programs. For Fig. 1A, time-course imaging began when colonies had been grown for 48hrs.

### Pulsed Field Gel Electrophoresis

PFGE protocols and analysis were conducted as described previously^58,59^.

### Statistical Analysis

To determine if specific chromosomal aneuploidies were enriched in one class of variant clones relative to the other (complex variants vs. smooth variants), we used a Chi square goodness of fit test and Pearson residuals analysis to compare the distribution of observed frequencies of CCNAs for each chromosome. Pearson residual values outside the range of -1.96 to +1.96 (two standard deviations) indicated significant enrichment of the specific chromosome between complex and smooth clones. From this, we determined that CCNAs of Chr13 and Chr15 displayed a significant enrichment in complex and smooth clones, respectively.

## Acknowledgements

We are grateful to Dr. Paul Magwene for sharing the strain YJM311. This study was supported by NIH/NIGMS awards 1K99GM13419301 to LRH and R35GM11978801 to JLA.

## Author Contributions

Conceptualization: LRH; Methodology: LRH; Investigation: LRH; Resources: LRH and JLA; Writing: LRH and JLA; Funding acquisition: JLA and LRH.

## Data and Materials Availability

Sequencing data for each clone in this study will be deposited on NCBI. All other data and materials will be made available upon request.

## Figure and Table Legends

**Supplemental Figure 1.**
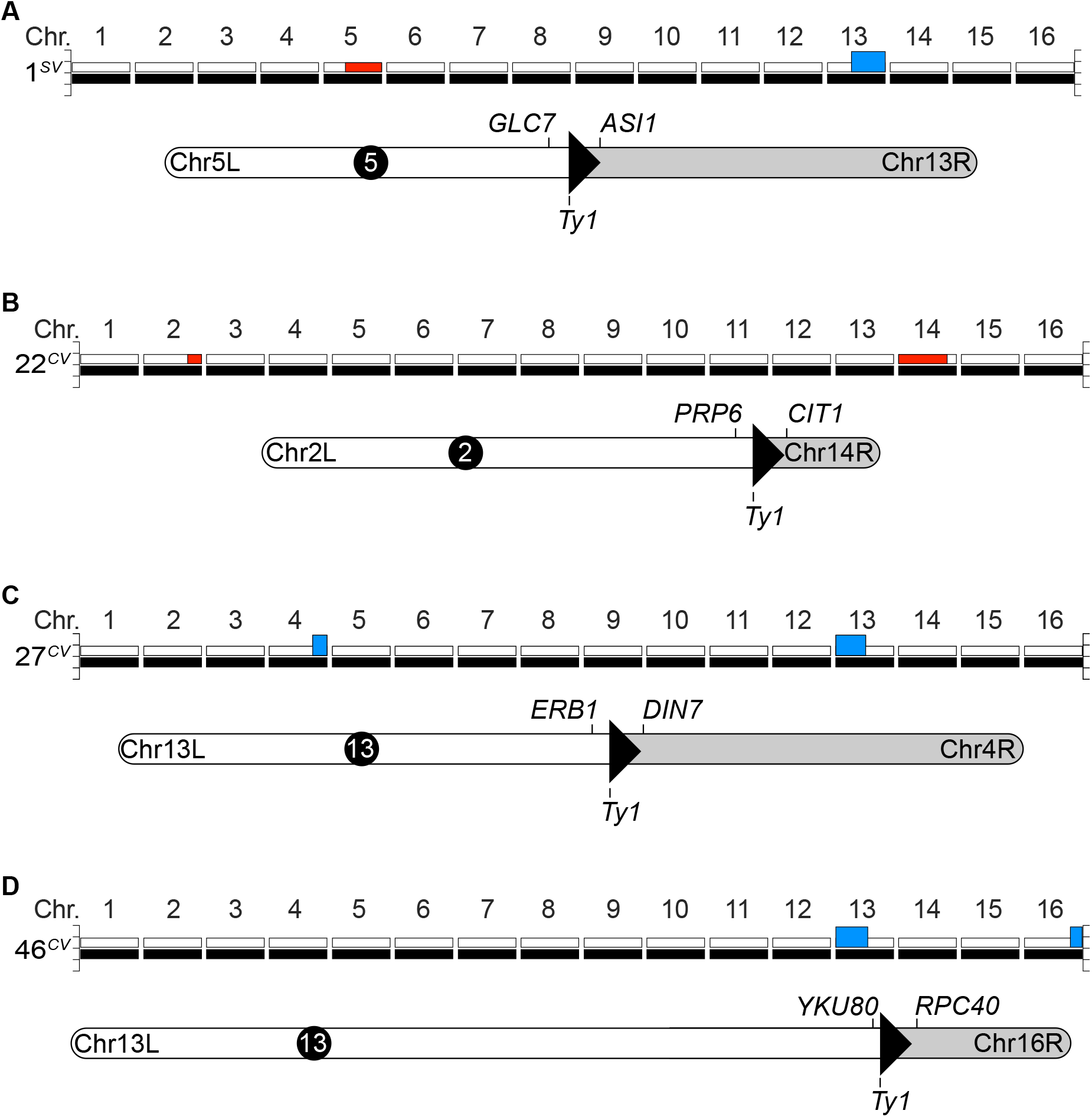
Structure of segmental CNVs identified in the genomes of phenotypic variants. **A-D**. Graphical karyotypes and cartoon schematics of translocations identified in the denoted variant genomes as deduced from Illumina and Oxford Nanopore sequencing analysis combined with pulsed field gel electrophoresis (PFGE). Detailed descriptions of these Seg. CNVs are presented in Table S2. For each chromosome, white bars represent homolog ‘*a*’ and black bars represent homolog ‘*b*’. Blue bars denote a copy number gain, red bars denote a copy number loss.

**Supplemental Figure 2.**
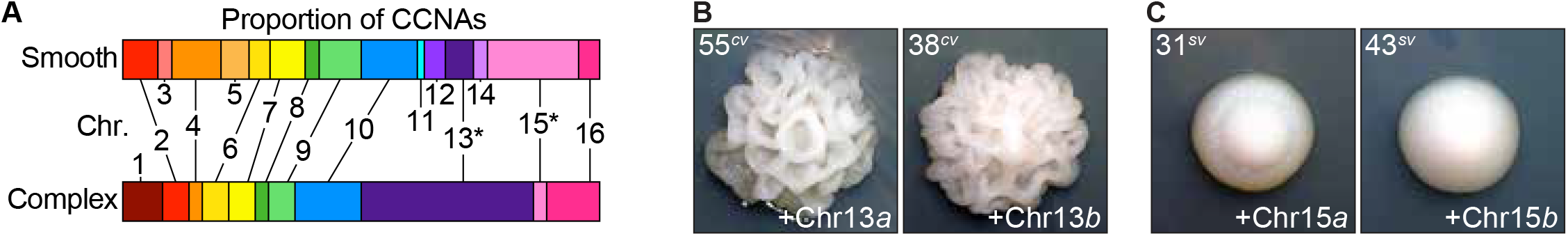
Copy number variations, not copy neutral changes in allelic identity, are dominant drivers of phenotypic variation. **A**. The proportion of total CCNAs that affected each chromosome in smooth and complex variants. Asterisk denotes an aneuploidy that is significantly enriched in one class of variant clones relative to the other. **B-C**. Representative images of colony morphologies displayed by the denoted variants. Colonies depicted in B. were grown on YP^2.0%^ plates. Colonies depicted in C. were grown on YP^0.5%^ plates.

**Supplemental Figure 3.**
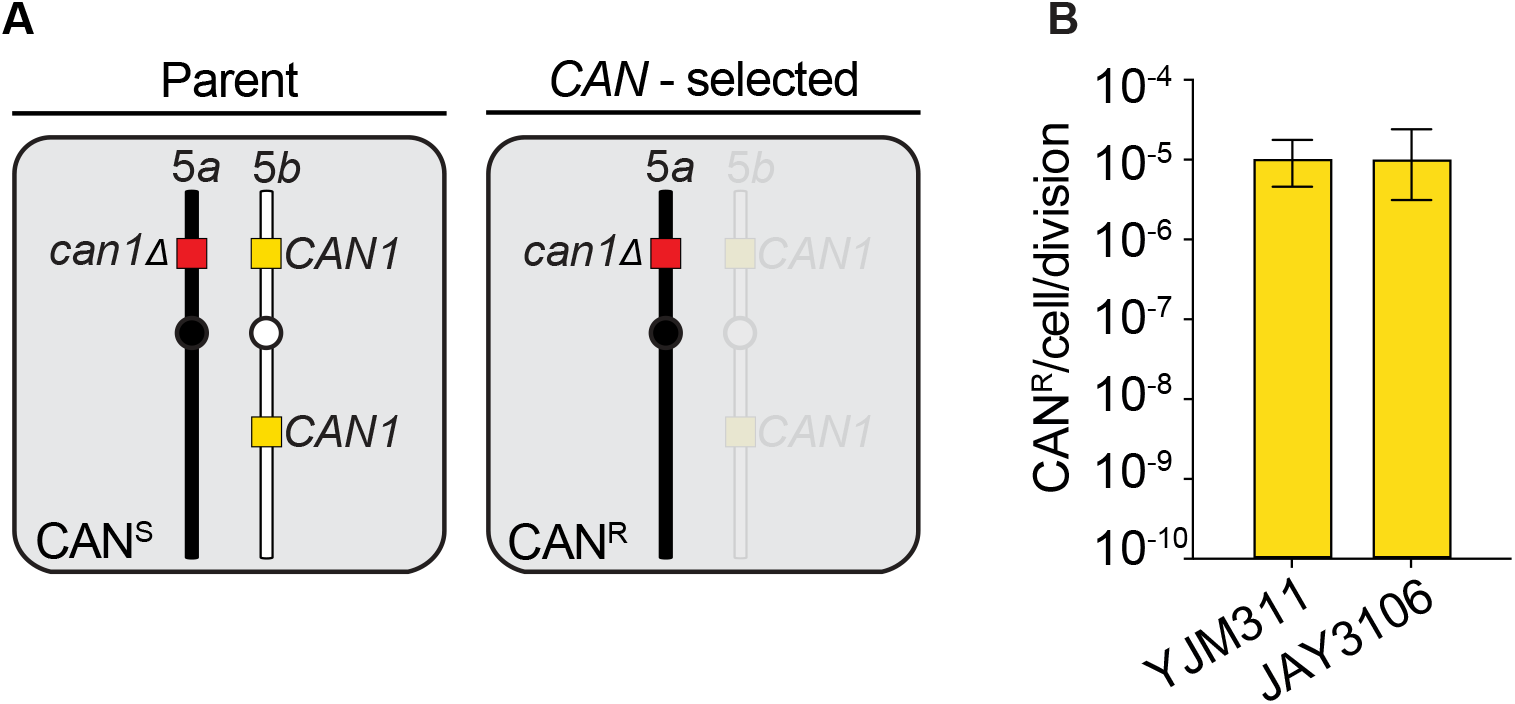
YJM311 does not express a CIN phenotype. **A**. A schematic illustrating the genotypic and phenotypic outcomes of the counter-selectable chromosome loss assay. *CAN1* deletion, red box; Chr5*b*-located *CAN1* markers, yellow boxes. **B**. The rates at which YJM311 and the laboratory diploid strain JAY3106 lost the *CAN1* marked homolog of Chr5. Error bars denote 95% confidence intervals.

**Supplemental Figure 4.**
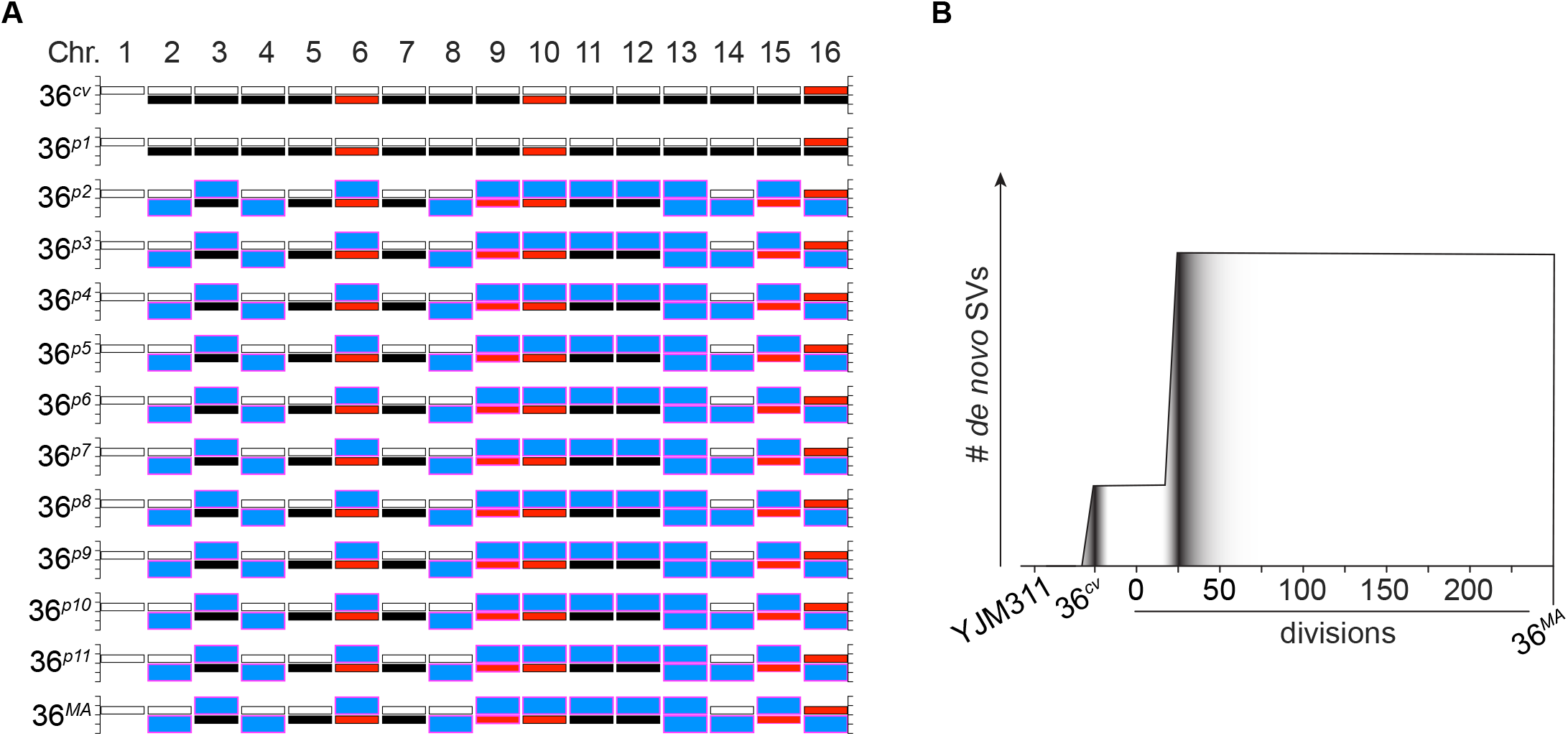
The karyotypic evolution of 36^*cv*^ over the course of the MA experiment. **A**. Graphical karyotypes of the 36^*cv*^ MA samples preserved at each progressive passage point. Bar colors same as in Fig. 3. **B**. A graphical representation depicting the genome evolution dynamics of 36^*MA*^ over the duration of the MA experiment.

## Supplemental Tables

**Table S1.** Copy Number Analysis of Chr1 in YJM311 and WT derivative clones. For comparison, copy number analysis for denoted region of Chr7 is also shown. Blue cells highlight WT clones that are monosomic for Chr1.

|  | genomic region:<br>chr1:50,000-150,000; 427<br>hetSNPs |  | genomic region:<br>chr7:200,000-300,000; 66<br>hetSNPs |  |
| --- | --- | --- | --- | --- |
| Isolate | log2<br>coverage<br>(median) | Copy Number | log2<br>coverage<br>(median) | Copy Number |
| YJM311 | -0.18 | 1.76 | -0.02 | 1.97 |
| 57 | -0.07 | 1.91 | -0.02 | 1.97 |
| 58 | -0.03 | 1.96 | -0.01 | 1.98 |
| 59 | -0.24 | 1.69 | -0.02 | 1.97 |
| 60 | -0.04 | 1.95 | -0.03 | 1.95 |
| 61 | -0.07 | 1.91 | -0.03 | 1.96 |
| 62 | -0.08 | 1.89 | -0.02 | 1.97 |
| 63 | -0.85 | 1.11 | -0.02 | 1.97 |
| 64 | -0.88 | 1.09 | -0.03 | 1.95 |
| 65 | -0.04 | 1.94 | -0.01 | 1.98 |
| 66 | -0.84 | 1.11 | -0.04 | 1.95 |
| 77 | -0.09 | 1.88 | -0.03 | 1.96 |
| 78 | -0.12 | 1.84 | -0.01 | 1.98 |
| 79 | -0.04 | 1.94 | -0.02 | 1.97 |
| 80 | -0.08 | 1.89 | -0.02 | 1.97 |
| 81 | -0.88 | 1.09 | -0.03 | 1.96 |
| 82 | -0.17 | 1.77 | -0.02 | 1.98 |
| 83 | -0.05 | 1.93 | -0.01 | 1.98 |
| 84 | -0.05 | 1.93 | -0.03 | 1.96 |
| 85 | -0.14 | 1.82 | -0.01 | 1.98 |
| 86 | -0.05 | 1.93 | -0.01 | 1.99 |

**Table S2.**
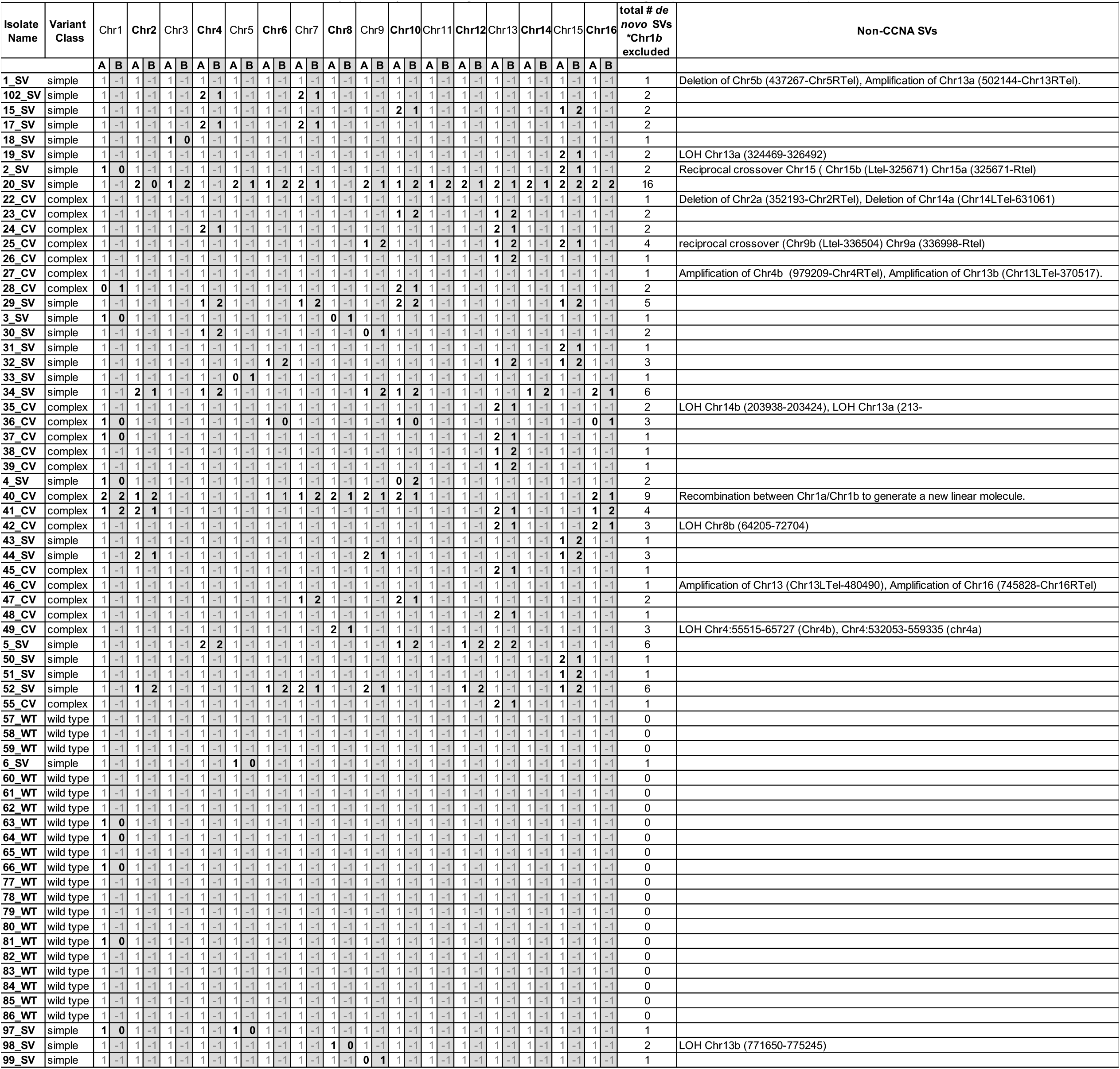
Karyotypic analysis of variant genomes. A/B delineate each homolog in a pair. Bold cells indicate aneuploidies.

**Table S3.** Fluctuation analysis derived rates of Chr5 loss in YJM311 background

|  |  |  |  |  |
| --- | --- | --- | --- | --- |
| Selection | Rate | Upper 95% difference | Lower 95% difference | Number of cultures |
| YJM311 | 1.04E-05 | 7.43E-06 | 5.80E-06 | 24 |
| JAY3106 | 1.03E-05 | 1.39E-05 | 7.14E-06 | 12 |

**Table S4.**
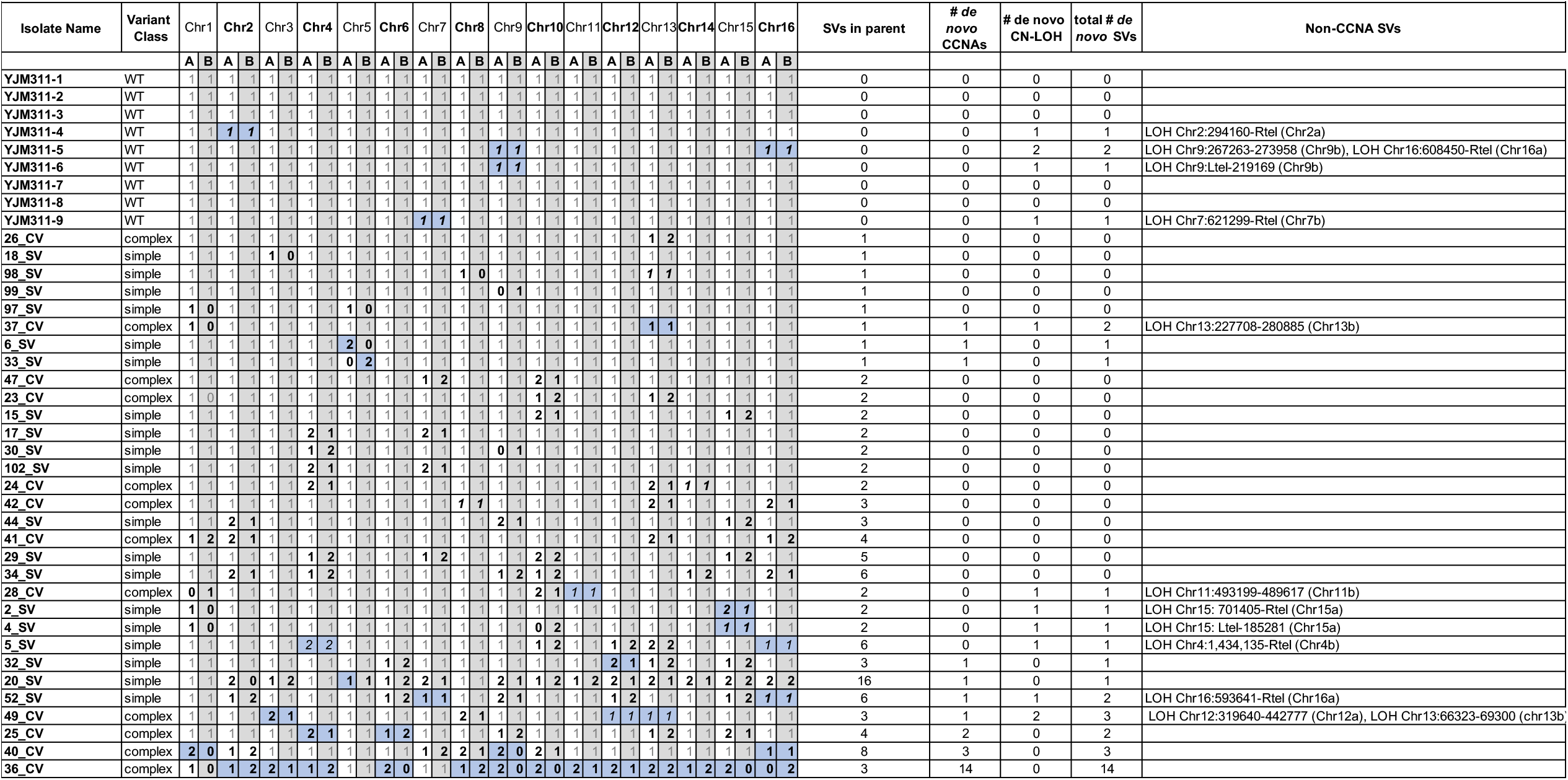
Karyotypic analysis of mutation accumulation lines derived from variant clones. A/B delineate each homolog in a pair. Bold cells indicate structural variations. Blue cells indicate de novo SVs gained during MA experiment.

